# Morphogenesis is transcriptionally coupled to neurogenesis during peripheral olfactory organ development

**DOI:** 10.1101/725705

**Authors:** Raphaël Aguillon, Romain Madelaine, Harendra Guturu, Sandra Link, Pascale Dufourcq, Virginie Lecaudey, Gill Bejerano, Patrick Blader, Julie Batut

**Affiliations:** Centre de Biologie du Développement (CBD, UMR5547), Centre de Biologie Intégrative (CBI, FR 3743), Université de Toulouse, CNRS, UPS, 31062, France; Department of Electrical Engineering, Stanford University, Stanford, CA 94305, USA; BIOSS Centre for Biological Signalling Studies, Albert Ludwigs University of Freiburg, Freiburg im Breisgau, Germany; Department of Developmental Biology, Department of Computer Science, Department of Pediatrics, Department of Biomedical Data Science, Stanford University, Stanford, CA 94305, USA

## Abstract

Sense organs acquire their distinctive shapes concomitantly with the differentiation of sensory cells and neurons necessary for their function. While our understanding of the mechanisms controlling morphogenesis and neurogenesis in these structures has grown, how these processes are coordinated remains largely unexplored. Neurogenesis in the zebrafish olfactory epithelium requires the bHLH proneural transcription factor Neurogenin1 (Neurog1). To address whether Neurog1 also controls morphogenesis in this system, we analysed the morphogenetic behaviour of early olfactory neural progenitors in *neurog1* mutant embryos. Our results indicate that the oriented movements of these progenitors are disrupted in this context. Morphogenesis is similarly affected by mutations in the chemokine receptor gene, *cxcr4b*, suggesting it is a potential Neurog1 target gene. We find that Neurog1 directly regulates *cxcr4b* through an E-boxes cluster located just upstream of the *cxcr4b* transcription start site. Our results suggest that proneural transcription factors, such as Neurog1, directly couple distinct aspects of nervous system development.

**One Sentence Summary:** Neurog1 controls olfactory organ morphogenesis via *cxcr4b*

## Introduction

The morphology of sense organs of the head is exquisitely adapted for detecting specific stimuli. At the same time that morphogenetic movements sculpt these structures during development, cell types are specified that will participate in their function either by detecting specific stimuli or transmitting sensory information to the brain. There is a growing literature concerning the molecular mechanisms controlling morphogenesis and specification of different cell types in sensory organs. Whether morphogenesis and cell fate specification are linked molecularly during the development of these organs, on the other hand, is unclear.

The zebrafish olfactory epithelium develops from a horseshoe-shaped pool of neural progenitors located at the boundary between the anterior neural plate and flanking non-neural ectoderm (Miyasaka et al., 2013). Neurogenesis in this system occurs in two distinct waves (Blader et al., 1997; Madelaine et al., 2011). Between 10 and 24 hours post-fertilisation (hpf), a set of early-born olfactory neurons (EON) differentiates. These neurons act as pioneers during the establishment of projections of olfactory sensory neurons (OSN) to the olfactory bulb, which are born during the second wave. Once OSN projections are established, a subset of EONs dies by apoptosis (Whitlock and Westerfield, 1998). The development of both EON and OSN require the partially redundant function of the bHLH proneural transcriptions factors Neurog1 and Neurod4 (Madelaine et al., 2011).

Concomitant with the earliest wave of neurogenesis in the developing olfactory epithelium, morphogenetic movements shape olfactory progenitors and newly born EON from territory into a placode (12-18 hpf) and then a rudimentary cup (18-24 hpf) (Whitlock and Westerfield, 2000; Breau et al., 2017). This process requires the chemokine receptor Cxcr4b, and its ligand Cxcl12a. Interfering with the activity of this signalling pathway, either by mis-expression of Cxcl12a or in *odysseus* (*ody*) embryos that carry mutations in *cxcr4b*, affects olfactory placode morphogenesis (Miyasaka et al., 2007).

In parallel to its role in olfactory neurogenesis, Neurog1 is ideally placed to control the cell movements that underlie morphogenesis of the olfactory cup, thus coupling morphogenesis and neurogenesis. Consistent with this idea, we find that the early phase of morphogenesis is compromised in *neurog1* mutant embryos. We provide evidence that the underlying defect is a lack of *cxcr4b* expression, which is directly regulated by Neurog1. Thus, we have uncovered a parsimonious mechanism for coordinating multiple features of peripheral sensory organ development, which we propose has been conserved with other vertebrates.

## Results and Discussion

To address a potential role for Neurog1 in the morphogenesis of the peripheral olfactory organ, we analysed its formation by time-lapse imaging *neurog1* mutant or control embryos carrying a *Tg(−8.4neurog1:gfp)* transgene (Golling et al., 2002; Blader et al., 2003); this transgene recapitulates the expression of endogenous *neurog1* during the development of the olfactory epithelium and has already been used as a short-term lineage label for the progenitors of early olfactory neurons, referred to hereafter as EON (Madelaine et al., 2011; Breau et al., 2017). As recently described by Breau and colleagues, we found that EON reach their final position in control embryos by converging towards a point close to the centre of the future cup (as represented in Figure 1A; (Breau et al., 2017)). Considering overall antero-posterior (AP) length of the EON population, this convergence appears to happen quickly until the olfactory placode has formed (12-18 hpf), after which it slows (Figure S1A,B and Movie S1). In *neurog1*^*hi1059*^ mutants, we observed a delay in convergence, which translates into a longer AP length spread of EON than seen in control embryos (Figure S1B and Movie S2). This delay is overcome, however, with the olfactory cup in *neurog1*^*hi1059*^ mutant embryos ultimately attaining AP length of control embryos (Figure S1B).

**Figure 1.**
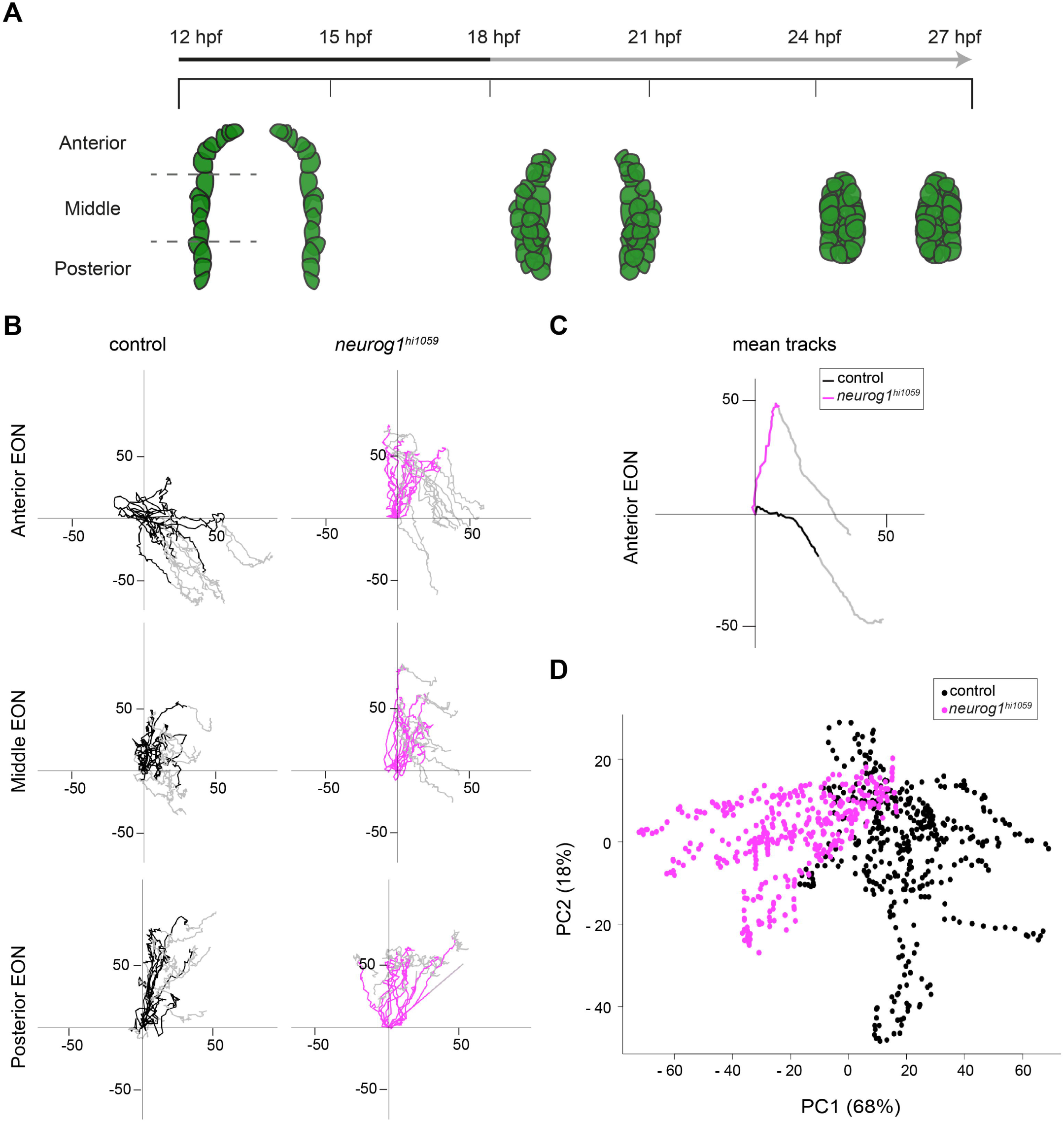
Oriented cell movements are affected in *neurog1*^*hi1059*^ mutant embryos during olfactory cup formation. (**A**) Graphic representation of the morphogenesis of olfactory cup from 12 hpf to 27 hpf showing a dorsal view of the three olfactory stages: olfactory territory (12 hpf), olfactory placode (18 hpf) and olfactory cup (24 hpf). EON progenitors are represented in green as visualised with the *Tg(−8.4neurog1:gfp)* transgene. At 12 hpf, the −8.4neurog1:GFP+ placodal domain can be divided in anterior, middle and posterior regions. The early (12-18 hpf; Black) and late (18-27 hpf; Grey) phases of morphogenesis are noted in the time line. (**B**) Tracks showing migration of EON of control (Black) or *neurog1*^*hi1059*^ mutant (Magenta) embryos. 12 tracks are represented (2 cells each from the left and right of 3 embryos) for each of the anterior, middle and posterior domains of the developing cup indicated in (**A**). The origin of the tracks has been arbitrarily set to the intersection of the X/Y axis and the early (coloured) and late (Grey) phases of migration have been highlighted. (**C**) Mean tracks for anterior EON of control (Black) or *neurog1*^*hi1059*^ mutant (Magenta) embryos. (**D**) Pairwise principal component analysis scatterplots of morphogenetic parameters extracted from the datasets corresponding to the tracks in (**C**). The major difference between control and *neurog1*^*hi1059*^ mutant embryos (PC1) corresponds to the antero-posterior axis.

To assess the morphogenetic phenotype of *neurog1*^*hi1059*^ mutant embryos at cellular resolution, we injected synthetic mRNAs encoding Histone2B-RFP (H2B-RFP) into *Tg(−8.4neurog1:gfp)* transgenic embryos, which were again imaged from 12 to 27 hpf. Morphogenetic parameters of individual EON located in the anterior, middle and posterior thirds of the initial neurog1:GFP+ population were extracted from datasets generated by manually tracking H2B-RFP positive nuclei (Movie S3 and S4). The position of each tracked EON was then plotted relative to its origin. As for the global analysis, the behaviour we observe for single EON in control embryos largely recapitulates those already reported (Figure 1B; (Breau et al., 2017)). Comparing the behaviour of EON in *neurog1*^*hi1059*^ mutants and siblings, we found that whereas EON in the middle and posterior regions of *neurog1*^*hi1059*^ mutant embryos migrate similarly to control siblings, the migratory behaviour of anterior EON is profoundly affected from 12 to 18 hpf (Figure 1B,C and Figure S2A); morphogenetic movements of individual skin cells showed no obvious differences in control versus *neurog1*^*hi1059*^ mutants suggesting that the effect is specific to EON (Figure S3). Principal component analysis (PCA) of the morphometric datasets confirmed that the major difference between control and *neurog1*^*hi1059*^ mutant embryos (PC1) lies in the migratory behaviour of anterior EON along the AP axis (Figure 1D); PCA revealed a more subtle difference in migration of the middle EON population along the same axis (Figure S2B) and between the posterior EON populations along the superficial-deep axis (Figure S2B). These migratory defects are not due to a decrease in cell mobility as EON in *neurog1* mutants displayed increased displacement over time compare to controls (Figure S1C,D); little or no difference was detected in the displacement of skin cells between control and *neurog1*^*hi1059*^ mutant embryos (Figure S1C,E). Taken together, our results indicate that Neurog1 is required between 12 and 18 hpf for the morphogenetic behaviour of olfactory progenitors.

The chemokine receptor Cxcr4b and its ligand Cxcl12a have been implicated in olfactory cup morphogenesis in the zebrafish (Miyasaka et al., 2007). To address whether the behaviour of EON in *neurog1*^*hi1059*^ mutants resembles that caused when the activity of this guidance receptor/ligand pair is abrogated, we analysed the morphogenetic parameters of EON in *cxcr4b*^*t26035*^ and *cxcl12a*^*t30516*^ mutants (Knaut et al., 2003; Valentin et al., 2007). As previously reported, olfactory progenitors in embryos lacking Cxcr4b or Cxcl12a function display convergence defects, highlighted by an increase in the AP length relative to controls (Figure S4A and Movies S5 and S6; (Miyasaka et al., 2007)). Analysis of the behaviour of individual EON in *cxcr4b*^*t26035*^ and *cxcl12a*^*t30516*^ mutant embryos indicates that defects in their migration are largely restricted to the anterior cohort (Figure 2A,B and Figure S5A,B); EON show increased displacement over time in both *cxcr4b*^*t26035*^ and *cxcl12a*^*t30516*^ mutants (Figure S4B,C) and no difference is apparent in the behaviour of skin cells in either mutant relative to control siblings (Figure S4B,D and S6). A combined PCA of morphometric datasets for anterior EON of *neurog1*^*hi1059*^, *cxcr4b*^*t26035*^ and *cxcl12a*^*t30516*^ mutants confirms that the major difference in EON behaviour lies in their displacement along the AP axis (PC1; Figure 2C). Finally, unsupervised clustering of the PCA analysis reveals that there is more resemblance in the behaviour of anterior EON between the three mutants than between any single mutant and controls (Figure 2D).

**Figure 2.**
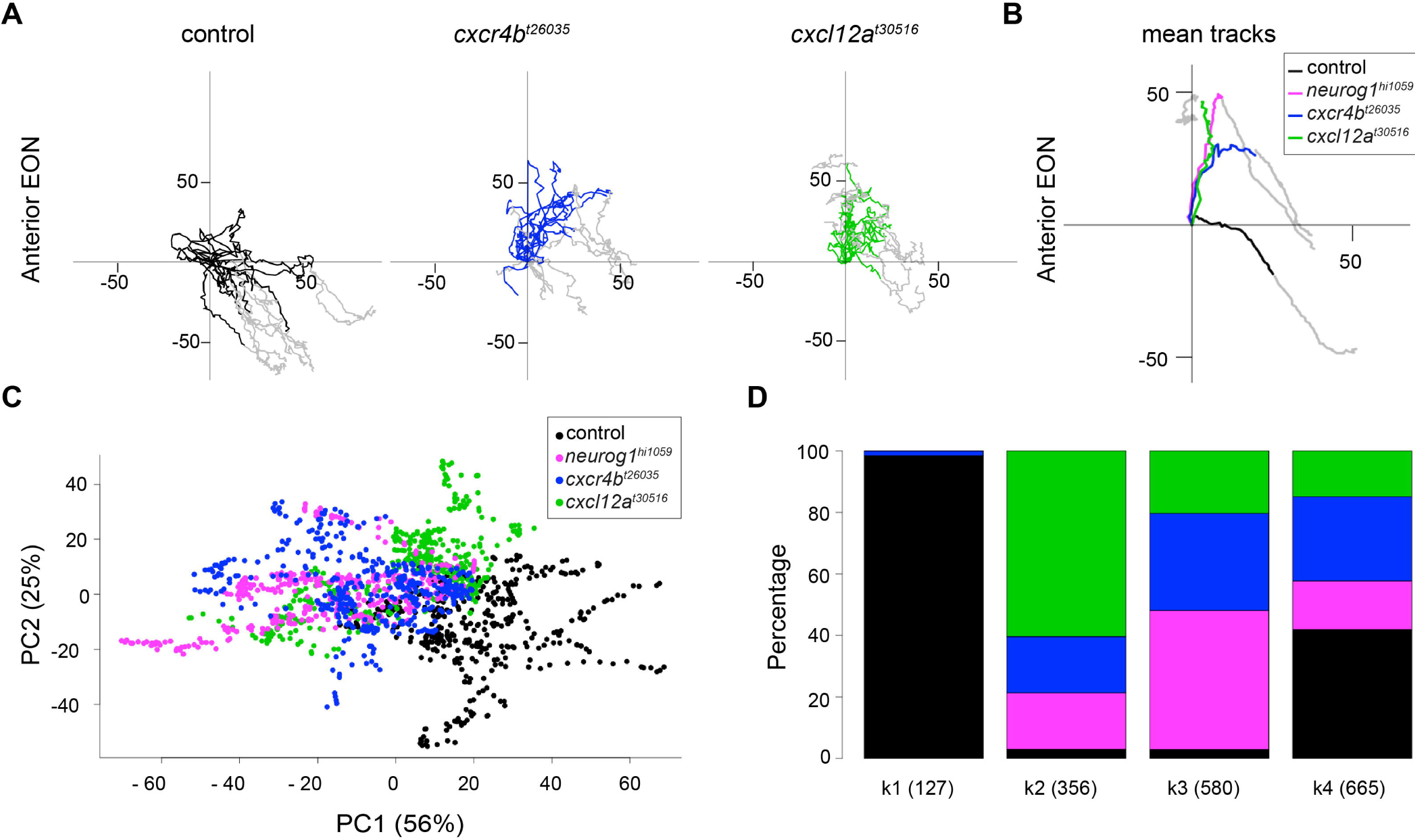
Morphogenetic defects in *cxcr4b*^*t26035*^ and *cxcl12a*^*t30516*^ mutant embryos resemble those of *neurog1*^*hi1059*^. (**A**) Tracks showing migration of anterior EON from control (Black), *cxcr4b*^*t26035*^ (Blue) and *cxcl12a*^*t30516*^ (Green) embryos. 12 anterior tracks are represented (2 cells each from the left and right of 3 embryos). The origin of the tracks has been arbitrarily set to the intersection of the X/Y axis and the early (coloured) and late (Grey) phases of migration have been highlighted. (**B**) Mean tracks showing migration of anterior EON of control (Black), *neurog1*^*hi1059*^ (Magenta), *cxcr4b*^*t26035*^ (Blue) and *cxcl12a*^*t30516*^ (Green) mutant embryos. (**C**) Pairwise principal component analysis scatterplots of morphogenetic parameters extracted from the datasets corresponding the tracks in (**B**). The major difference between control, *neurog1*^*hi1059*^, *cxcr4b*^*t26035*^ and *cxcl12a*^*t30516*^ mutant embryos (PC1) corresponds to the antero-posterior axis. (**D**) Clustering analysis of morphogenetic parameters extracted from the datasets presented in (**B**) and analysed in (**C**). One cluster, k1, contains almost exclusively control cells (Black), whereas cells from *neurog1*^*hi1059*^ (Magenta), *cxcr4b*^*t26035*^ (Blue) and *cxcl12a*^*t30516*^ (Green) mutant embryos clustered together in k2, k3 and k4.

The similarity in the migration phenotype of EON in *neurog1*^*hi1059*^, *cxcr4b*^*t26035*^ and *cxcl12a*^*t30516*^ mutant embryos suggests that the proneural transcription factor and the receptor/ligand couple act in the same pathway. To determine if the expression of either the receptor or its ligand are affected in the absence of Neurog1, we assessed their expression in *neurog1*^*hi1059*^ mutant embryos. We found that *cxcr4b* expression is dramatically reduced or absent in EON progenitors at 12 and 15 hpf in this context (Figure 3A); the expression of *cxcr4b* recovers in *neurog1*^*hi1059*^ mutant embryos from 18 hpf, a stage at which we have previously reported that the expression of a second bHLH proneural gene, *neurod4*, also becomes Neurog1-independent (Figure 3A; (Madelaine et al., 2011). Contrary to *cxcr4b*, the expression of *cxcl12a* is unaffected in *neurog1*^*hi1059*^ mutant embryos at all stages analysed (Figure 3B). Taken together, these results suggest that the EON migration phenotype in *neurog1*^*hi1059*^ mutant embryos results from the lack of Cxcr4b during the early phase of olfactory cup morphogenesis.

**Figure 3.**
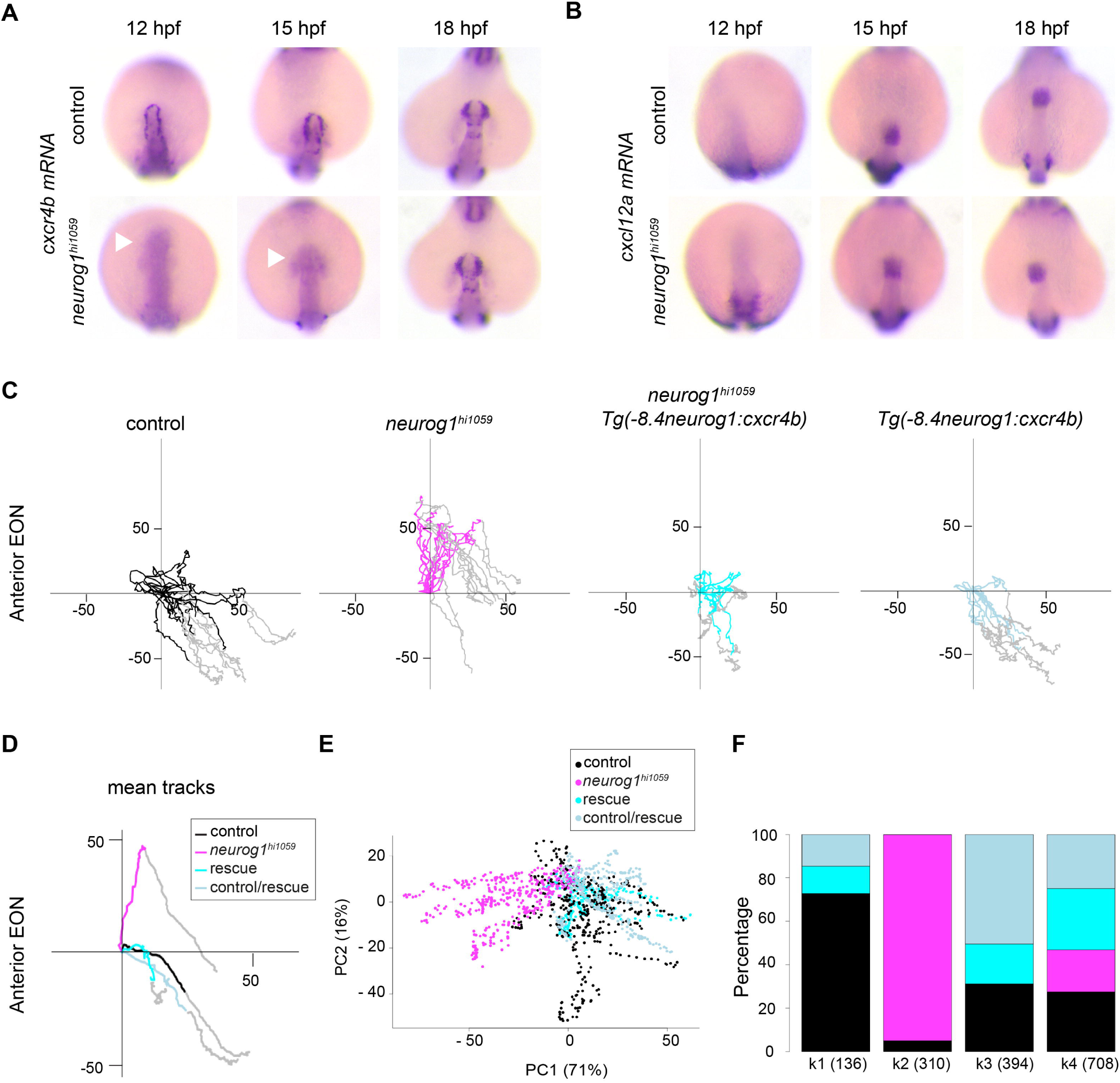
Cxcr4b is the predominant downstream effector of Neurog1 during olfactory cup morphogenesis. (**A**) *cxcr4b* whole mount *in situ* hybridisation at 12, 15 and 18 hpf in control and *neurog1*^*hi1059*^ mutant embryos. *cxcr4b* expression is dramatically reduced or absent in EON progenitors at 12 and 15 hpf in *neurog1*^*hi1059*^ mutant embryos (white arrowheads) but from 18 hpf the expression of *cxcr4b* recovers. (**B**) *cxcl12a* whole mount *in situ* hybridisation at 12, 15 and 18 hpf in control and *neurog1*^*hi1059*^ mutant embryos, in which *cxcl12a* expression is not affected. (**C**) Tracks showing migration of anterior EON of control (Black) embryos, *neurog1*^*hi1059*^ mutant embryos (Magenta), *neurog1*^*hi1059*^ mutant embryos carrying the *Tg(− 8.4neurog1:cxcr4b)* transgene (Cyan) and control embryos carrying *Tg(− 8.4neurog1:cxcr4b)* (Light Blue). 12 anterior tracks are represented (from 4 embryos). The origin of the tracks has been arbitrarily set to the intersection of the X/Y axis and the early (coloured) and late (Grey) phases of migration have been highlighted. (**D**) Mean tracks showing migration of anterior EON of the tracks in (**C**). (**E**) Pairwise principal component analysis scatterplots of morphogenetic parameters extracted from the datasets presented in (**D**). The major difference between control, *neurog1*^*hi1059*^, and control or *neurog1*^*hi1059*^ with the rescue transgene (PC1) corresponds to the antero-posterior axis. (**F**) Clustering analysis of morphogenetic parameters extracted from the datasets presented in (**D**) and analysed in (**E**). One cluster, k2, contains only *neurog1*^*hi1059*^ cells (Magenta), whereas cells from control (Black), rescue (Cyan) and control/rescue (Light Blue) embryos clustered together in k1, k3 and k4.

If the absence of early *cxcr4b* expression in *neurog1*^*hi1059*^ mutants underlies the morphogenesis defects in this background, we hypothesised that restoring its expression should rescue these defects. To test this, we generated a transgenic line where expression of the chemokine receptor is controlled by a −8.4 kb fragment of genomic DNA responsible for *neurog1* expression in EON, *Tg(−8.4neurog1:cxcr4b- mCherry)*, and introduced it into the *neurog1*^*hi1059*^ mutant background (Blader et al., 2003; Madelaine et al., 2011). Analysis of the migratory behaviour of anterior EON in *neurog1*^*hi1059*^ mutant embryos carrying the transgene indicates that they display oriented posterior migration similar to control embryos and siblings carrying the transgene (Figure 3C,D and Movies S7 and S8). The similarity in the behaviour of the anterior EON is also evident after PCA analysis, where *neurog1*^*hi1059*^ mutant cells carrying the transgene cluster primarily with control cells with or without the transgene rather than mutant cells lacking the transgene (Figure 3E,F). These data lead us to conclude that Cxcr4b is the predominant downstream effector of Neurog1 during the early phase of olfactory cup morphogenesis.

Finally, we asked whether *cxcr4b* is a direct transcriptional target of Neurog1 by searching for potential Neurog1-dependent cis-regulatory modules (CRM) at the *cxcr4b* locus. Proneural transcription factors bind CANNTG sequences known as E-boxes, which are often found in clusters (Bertrand et al., 2002). We identified 18 E-boxes clusters in the sequences from −100 to +100 kb of the *cxcr4b* initiation codon, but only 1 of these clusters contains more than one of the CA^A^/_G_ATG E-box sequence preferred by Neurog1 (Figure 4A and data not shown; (Madelaine and Blader, 2011)). Coherent with a role for this E-box cluster in the regulation of *cxcr4b* expression, a transgenic line generated using a 35kb fosmid clone that contains this cluster, *TgFOS(cxcr4b:eGFP)*, shows robust expression of GFP in the olfactory cup (Figure 4A,D). To investigate whether this cluster acts as a *bona fide* Neurog1-dependent CRM, we performed chromatin immunoprecipitation (ChIP) experiments. In the absence of a ChIP compatible antibody against endogenous zebrafish Neurog1, we chose a strategy based on mis-expression of a Ty1-tagged form of Neurog1. Mis-expression of Neurog1- Ty1 efficiently induces the expression of *deltaA*, a known Neurog1 target, and *cxcr4b* suggesting that tagging Neurog1 does not affect its transcriptional activity and that *cxcr4b* behaves as a Neurog1 target (Figure 4B). We have previously shown that the *deltaA* locus contains two proneural regulated CRMs (Madelaine and Blader, 2011); whereas CRM HI is Neurog1-dependent, HII underlies regulation of *deltaA* by members of the Ascl1 family of bHLH proneural factors (Hans and Campos-Ortega, 2002; Madelaine and Blader, 2011). We found that ChIP against Neurog1-Ty1 after mis-expression effectively discriminates between the Neurog1-regulated HI and Ascl1- regulated HII CRM at the *deltaA* locus, thus providing a control for the specificity of our ChIP strategy (Figure 4C). Similarly, we were able to ChIP the potential CRM containing the CAGATG E-box cluster at the *cxcr4b* locus suggesting that this region is also a target for Neurog1 (Figure 4C).

**Figure 4.**
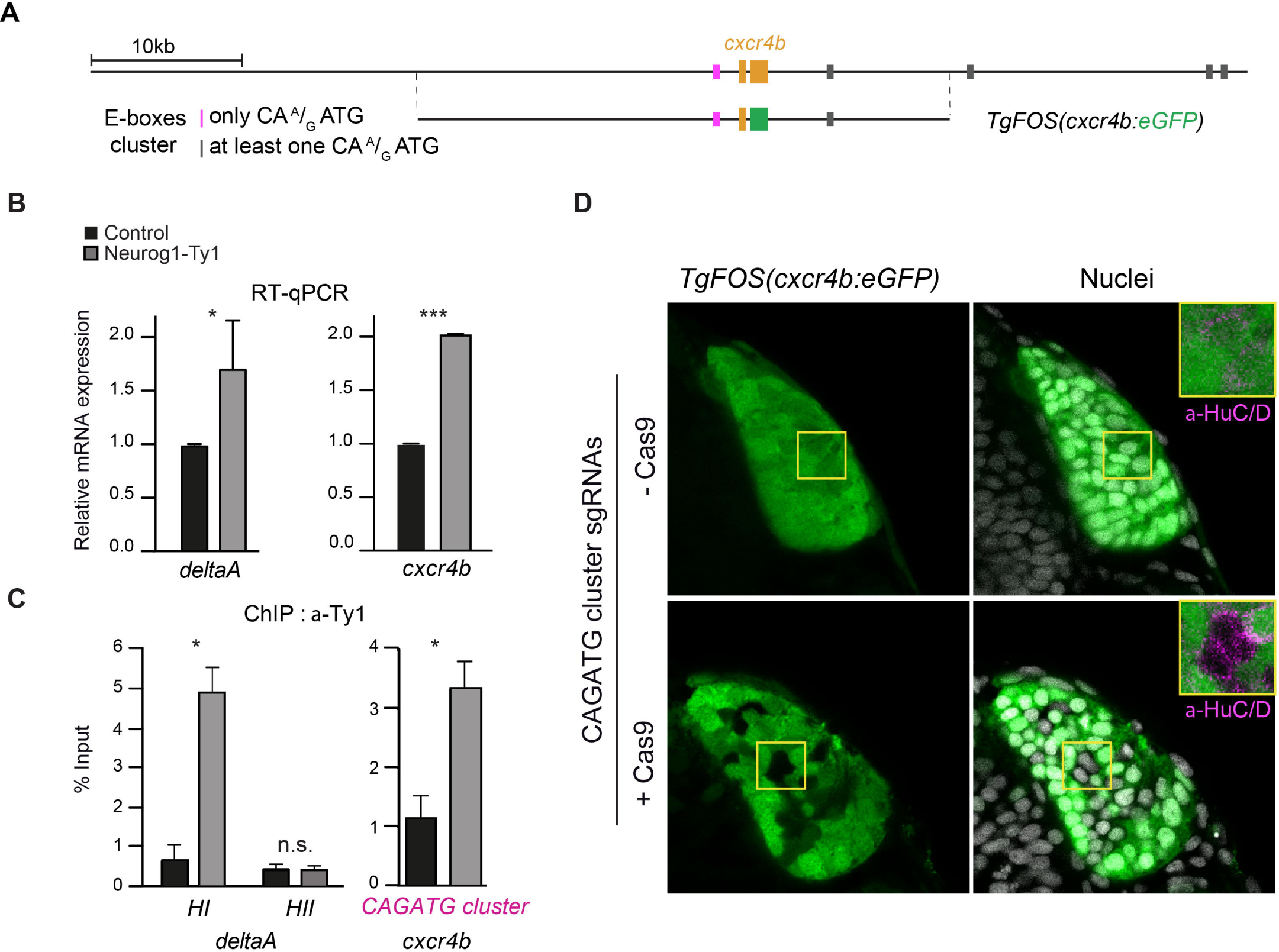
Neurog1 directly controls *cxcr4b* expression via an upstream Cis-regulatory module (CRM). (**A**) Schematic representation of the *cxcr4b* locus indicating the position of exons of the *cxcr4b* gene (orange) and E-box clusters, which are color-coded depending on the nature of the E-box sequences. Also presented is the position of the genomic sequences found in the *TgFOS(cxcr4b:eGFP)* transgene. (**B**) qPCR analysis of the effect of Neurog1-Ty1mRNA mis-expression on the relative mRNA levels of the known Neurog1 target gene *deltaA* and *cxcr4b*. A significant increase in expression is detected for both genes. Shown are mean ± s.e.m, p values are calculated using a two-tailed Student’s t-test, *p=0.01, ***p=0.0001. (**C**) Chromatin immunoprecipitation (ChIP) using an antibody against Ty1 and chromatin prepared from 15 hpf embryos mis-expressing Neurog1-Ty1 mRNA (Grey). Control (Black) represents ChIP with IgG alone. Shown are mean ± s.e.m, p values are calculated using a two-tailed Student’s t-test, n.s. not significant, *p=0.01. (**D**) Single confocal sections of *TgFOS(cxcr4b:eGFP)* embryos at 24 hpf showing eGFP expression in the olfactory cups, and either HuC/D expression or nuclear labelling (TOPRO). Embryos were injected with an sgRNA pair flanking the E-box containing CRM at the *cxcr4b* locus plus or minus Cas9 as a control. Insets show HuC/D expression in both conditions.

To address the importance of the E-box cluster in the regulation of *cxcr4b* expression, we employed a Crispr/Cas9 approach to delete this CRM using a pair of sgRNAs flanking the CRM (Figure S7A). The sgRNA pair efficiently induces deletions in the targeted sequence, as judged by PCR on genomic DNA extracted from injected embryos (Figure S7B). Injection of the sgRNA pair into *TgFOS(cxcr4b:eGFP)* transgenic embryos caused mosaic disruption of the eGFP expression pattern (Figure 4D). Loss of *TgFOS(cxcr4b:eGFP)* transgene expression is not due to cell death as eGFP-negative cells maintain the expression of the early neuronal marker HuC/D (inserts in Figure 4D). Taken together, the results from our ChIP and Cripsr/Cas9 experiments strongly suggest that the CAGATG E-box cluster upstream of *cxcr4b* is regulated directly by Neurog1.

Neurog1 controls an early wave of neurogenesis in the zebrafish olfactory epithelium (Madelaine et al., 2011). As in invertebrates, control of neurogenesis by this proneural transcription factor is achieved via the transcriptional regulation of so-called neurogenic genes, such as *deltaA* and *deltaD* in the fish (Hans and Campos-Ortega, 2002; Madelaine and Blader, 2011). Our present study highlights that Neurog1 is also required for morphogenesis of the zebrafish peripheral olfactory sensory organ, in this case via its target gene *cxcr4b*. Thus, our data supports a simple mechanism whereby Neurog1 couples neurogenesis with morphogenesis via the transcriptional regulation of distinct targets. It has previously been shown that members of the Neurog family regulate *Delta1* and *Cxcr4* expression in the mouse, and that development the olfactory epithelium in this model requires Neurog-family proneural factors (Beckers et al., 2000; Mattar et al., 2004; Shaker et al., 2012). We propose, therefore, that this simple mechanism for coordinating the development of the olfactory system has been conserved across vertebrates. More generally, we can envisage that the role of proneural factors is retained in both neuronal fate and morphogenesis instruction and we can stipulate that the specific shape of each tissue and/or organ is also dictated by a combination of bHLH factors.

## Acknowledgments

This work was supported by the Centre National de la Recherche Scientifique (CNRS); the Institut National de la Santé et de la Recherche Médicale (INSERM); Université de Toulouse III (UPS); Fondation pour la Recherche Médicale (FRM; DEQ20131029166); Fédération pour la Recherche sur le Cerveau (FRC); and the Ministère de la Recherche. We thank Kristen Kwan and Chi-Bin Chien for providing plasmids of the Tol2kit, Stéphanie Bosch, Brice Ronsin and the Toulouse LITC RIO Imaging platform, and Aurore Laire and Richard Brimicombe for taking care of the fish. We also thank Marie Breau, Magali Suzanne, Christian Mosimann and members of the Blader lab for advice on experiments and comments on the manuscript.

## Materials and Methods

### Fish Husbandry and lines

Ethics Statement and Embryos: All embryos were handled according to relevant national and international guidelines. French veterinary services and the local ethical committee approved the protocols used in this study, with approval ID: A-31-555-01 and APAPHIS #3653-2016011512005922v6.

Fish were maintained at the CBD-CBI zebrafish facility in accordance with the rules and protocols in place. The *neurog1*^*hi1059*^, *cxcr4b*^*t26035*^ and *cxcl12a*^*t30516*^ mutant lines have previously been described (Golling et al., 2002; Knaut et al., 2003; Valentin et al., 2007), as has the *Tg(−8.4neurog1:gfp)*^*sb1*^ (Blader et al., 2003). Embryos were obtained through natural crosses and staged according to (Kimmel et al., 1995).

### Establishment of new transgenic lines

The *Tg(−8.4neurog1:cxcr4b-mCherry)* transgene was generated by first cloning the coding region of *cxcr4b* minus its endogenous stop codon in frame upstream of mCherry. The resulting *cxcr4b-mCherry* fusion coding sequence was transferred into the middle entry plasmid of the Tol2kit developed in the Chien lab (Kwan et al., 2007). The final transgene vector was generated using LR recombination with a previously described *p5’-8.4neurog1* (Madelaine et al., 2011), the *pME-cxcr4b-mCherry*, and the *p3E-polyA* and *pDestTol2pA/pDestTol2pA2* from the Tol2kit (Kwan et al., 2007). The line was then generated by co-injecting the transgene with mRNA encoding Tol2 transposase into freshly fertilised zebrafish embryos.

The *TgFOS(cxcr4b:eGFP)*^*fu10Tg*^ transgenic line was generated using homologous recombination by replacing the second exon of *cxcr4b* by LynGFP in the Fosmid CH1073-406F3, followed by zebrafish transgenesis (Revenu et al., 2014). The first 5 amino acid encoded by the first exon of *cxcr4b* are fused to LynGFP, preventing targeting to the membrane. The GFP localises to the cytoplasm in this transgenic line.

### In situ Hybridisation, Immunostaining and Microscopy

In situ hybridisation was performed as previously described (Oxtoby and Jowett, 1993). Antisense DIG-labelled probes for *cxcr4b* and *cxcl12a* (David et al., 2002) were generated using standard procedures. In situ hybridisations were visualised using BCIP and NBT (Roche) as substrates.

Embryos were immunostained as previously described (Madelaine et al., 2011); primary antibody used was mouse anti-HuC/D (1:500; 16A11, Molecular Probes, USA), which was detected using Alexa Fluor 555 conjugated goat anti-mouse IgG diluted 1/1000: (A-28180, Molecular Probes, USA). Immunostained embryos were counterstained with Topro3 (T3605, Molecular Probes, USA). Labelled embryos were imaged using an upright SP8 Leica confocal and analysed using ImageJ and Imaris 8.3 (Bitplane, Switzerland) software.

### Cell tracking in time-lapse confocal datasets

Embryos carrying the *Tg(−8.4neurog1:gfp)* transgene (Blader et al., 2003) were injected with synthetic mRNA encoding an H2B-RFP fusion protein; for analysis of the global behaviour of olfactory morphogenesis, un-injected embryos were used. Embryos were then grown to 12 hpf at which point they were dechorionated and embedded for imaging in 0.7% low-melting point agarose in embryos medium. A time-lapse series of confocal stacks (1 mm slice/180 mm deep) was generated of the anterior neural plate and flanking non-neural ectoderm on an upright Leica SP8 Confocal microscope using a 25x HC Fluotar water-immersion objective. Acquisitions each 7 min were stopped at 27 hpf, when the olfactory rosette was clearly visible. The lineage of anterior, midline and posterior early olfactory neuron cohorts were subsequently constructed semi-automatically following H2B-RFP of neurog1:GFP+ EON using Imaris 8.3 analysis software (Bitplane, Switzerland); unless mentioned, for each of three embryos, two anterior, middle and posterior cells from the left and right olfactory organs were tracked.

### Track analysis

Track parameters were extracted from Imaris as Excel files and analysed using a custom script generated in R (The R Project for Statistical Computing, www.r-project.org). First, tracks were rendered symmetric across the left-right axis for ease of interpretation. Tracks were then colour coded according to their genotype and to the phase of migration (early from 12-18 hpf; late from 18-27 hpf) and plotted. Finally, the mean for each set of tracks was generated using the “RowMeans” function and a plot was generated.

Principal component analysis (PCA) and clustering were performed using the built-in R function “prcomp” from the “FactoMineR” package and the “kmeans” function, from the “stats” package, respectively. Finally, the “barplot” function (“graphics” package) was used to represent either the EON cluster composition or the Skin cluster composition.

### Chromatin Immunoprecipitation and qPCR

ChIP experiments were performed as previously described using approximately 300 embryos (12 to 15 hpf) per immunoprecipitation (Wardle et al., 2006). Two to four separate ChIP experiments were carried out with corresponding independent batches of either control un-injected embryos or embryos injected with a synthetic mRNA encoding Neurog1-Ty1; ChIP-grade mouse anti-Ty1 (BB2; Sigma-Aldrich, USA) was used. Primers used for qPCR on ChIPs were:

*cxcr4b* CATATG cluster fw 5’- CTACATCTAAAAATTGAAAGA-3’

*cxcr4b* CATATG cluster rev 5’- CAAACCCAACACCCCTACTG-3’

*deltaA* HI fw 5’- GCGGAATGAACCACCAACTT-3’

*deltaA* HI rev 5’- GTGTGACTAAAGGTGTATGGGTG-3’

*deltaA* HII fw 5’- TATTGTGTGCAGGCGGAATA-3’

*deltaA* HII rev 5’-GTTTGAATGGGCTCCTGAGA-3’.

Reactions were carried out in triplicates on a MyIQ device (Bio-Rad). The specific signals were calculated as the ratio between the signals with the Ty1 antibody and beads alone, and were expressed as percentage of chromatin input.

For qPCR experiments, to determine expression levels of *cxcr4b* and *deltaA* after mis-expression of Neurog1-Ty1, total RNAs were extracted from 20 injected embryos at 15 hpf with the RNeasy Mini Kit (QIAGEN), and reverse-transcribed with the PrimeScript RT reagent kit (Ozyme) according to the supplier’s instructions. Q-PCR analyses were performed on MyIQ device (Bio-Rad) with the SsoFast EvaGreen Supermix (Bio-Rad), according to the manufacturer’s instructions. All experiments include a standard curve. Samples from embryos were normalised to the number of *ef1a* mRNA copies. Primers for qPCR to determine the expression levels of *cxcr4b* and *deltaA* after mis-expression of Neurog1-Ty1 normalised to the expression of *ef1a* were:

*cxcr4b* coding fw 5’- GCTGGCATATTTCCACTGCT-3’

*cxcr4b* coding rev 5’- AGTGCACTGGACGACTCTGA-3’

*deltaA coding* fw 5’- CGGGTTTACAGGCATGAACT-3’

*deltaA coding* rev 5’- ATTGTTCCTTTCGTGGCAAG-3’

*ef1a* fw 5’-GCATACATCAAGAAGATCGGC-3’

*ef1a* rev 5’-GCAGCCTTCTGTGCAGACTTTG-3’.

### Crispr/Cas9 deletion of potential CRM at the *cxcr4b* locus

sgRNA sequences flanking the CAGATG E-box cluster at the *cxcr4b* locus were designed using the web-based CRIPSRscan algorithm (Moreno-Mateos et al., 2015); http://www.crisprscan.org). The targeted sequences are 5’- GGCTTATGATGGAGGCGAC<u>TGG</u>-3’ and 5’-GGCTTGTATTGCCCTTGA<u>GGG</u>-3’; the PAM sequence at the target site are underlined. Templates for the transcription of sgRNAs were generated by PCR following previously described protocols (Nakayama et al., 2014). Injection of sgRNAs was performed as described by Burger and colleagues, using a commercially available Cas9 protein (New England Biolabs). The efficiency of creating deletion after co-injection of the sgRNA pair was determined by PCR on genomic DNA extracted from injected embryos using the following primers:

5’-AACTCGCATTCGGCAAACTCTC-3’

5’-AAGGGGATAATGAGCAGTCAGC-3’.

While a 500 base-pair PCR fragment is generated from a wild-type locus, an approximately 200 base-pair fragment is amplified if a deletion has been induced.

## Supplemental Figures

**Figure S1.** *neurog1* mutant embryos display specific defects in olfactory development.

(**A**) Schematic representation of the relative antero-posterior length calculation. Lengths are normalised relative to the 12 hpf antero-posterior length.

(**B**) Graph showing normalised antero-posterior length of the developing olfactory cups in control (Black) and *neurog1*^*hi1059*^ mutant embryos (Magenta) over time. The mean ±;s.e.m of 3 embryos/6 cups are represented per condition.

(**C**) Histogram showing the sum of the displacement of EON (empty) and skin cells (checkered) of control (Black) and *neurog1*^*hi1059*^ mutant embryos (Magenta) along the antero-posterior axis during olfactory cup development. The mean ± s.e.m of 12 cells are represented per condition.

(**D**) Graph showing the sum of the displacement along the antero-posterior axis of control (Black) or *neurog1*^*hi1059*^ mutant (Magenta) EON during five indicated time periods. The mean ± s.e.m of 36 cells are represented (12 cells per embryo and 3 embryos per genotype) per condition.

(**E**) Graph showing the sum of the displacement along the antero-posterior axis of control (Black) or *neurog1*^*hi1059*^ mutant (Magenta) skin cells during five indicated time periods. The mean ± s.e.m of 36 cells are represented (12 cells per embryo and 3 embryos per genotype) per condition.

**Figure S2.** Anterior EON population tracks are specifically affected in *neurog1* mutated embryos.

(**A**) Mean tracks for anterior, middle and posterior EON of control (Black) or *neurog1*^*hi1059*^ mutant (Magenta) embryos. Means are from 12 tracks (2 cells each from the left and right of 3 embryos) for each region. The origin of the tracks has been arbitrarily set to the intersection of the X/Y axis and the early (coloured) and late (Grey) phases of migration have been highlighted.

(**B**) Pairwise principal component analysis scatterplots of morphogenetic parameters extracted from the datasets presented in (**A**). Whereas for anterior PC1 and PC2 respectively correspond to the antero-posterior and medio-lateral axes, for middle and posterior PC1 and PC2 represent the antero-posterior and superficial-deep axes.

**Figure S3.** Morphogenetic movements of skin cells in *neurog1*^*hi1059*^ mutant embryos. Tracks showing migration of skin cells in control (Black) or *neurog1*^*hi1059*^ mutant (Magenta) embryos. 12 tracks are represented for cells overlying the anterior, middle and posterior domains of the developing cup. The origin of the tracks has been arbitrarily set to the intersection of the X/Y axis and the early (coloured) and late (Grey) phases of migration have been highlighted. Morphogenetic movements of individual skin cells showed no obvious differences in control versus *neurog1* mutants.

**Figure S4.** Morphogenetic movements of EON are globally affected in *cxcr4b*^*t26035*^ and *cxcl12a*^*t30516*^ mutant embryos.

(**A**) Graph showing normalised antero-posterior length of the developing olfactory cup in control (Black) and *cxcr4b*^*t26035*^ (Blue) and *cxcl12a*^*t30516*^ (Green) mutants over time. The mean ± s.e.m of 3 embryos/6 cups are represented per condition.

(**B**) Histogram showing the sum of the displacement of EON (empty) and skin cells (checkered) for control (Black), *neurog1*^*hi1059*^ (Magenta) *cxcr4b*^*t26035*^ (Blue) and *cxcl12a*^*t30516*^ (Green) mutants along the antero-posterior axis during olfactory cup development. The mean ± s.e.m of 12 cells are represented per condition.

(**C**) Graph showing the sum of the displacement along the antero-posterior axis of control (Black), *neurog1*^*hi1059*^ (Magenta) *cxcr4b*^*t26035*^ (Blue) and *cxcl12a*^*t30516*^ (Green) EON during five indicated time periods. The mean ± s.e.m of 36 cells are represented (12 tracks per embryo and 3 embryos per genotype) per condition.

(**D**) Graph showing the sum of the displacement along the antero-posterior axis of control (Black), *neurog1*^*hi1059*^ (Magenta) *cxcr4b*^*t26035*^ (Blue) and *cxcl12a*^*t30516*^ (Green) skin cells during five indicated time periods. The mean ± s.e.m of 36 cells are represented (12 tracks per embryo and 3 embryos per genotype) per condition.

**Figure S5.** Morphogenetic parameters in *cxcr4b*^*t26035*^ and *cxcl12a*^*t30516*^ mutant embryos.

(**A**) Tracks showing migration of EON of control (Black), *cxcr4b*^*t26035*^ (Blue) and *cxcl12a*^*t30516*^ (Green) embryos. 12 tracks are represented (2 cells each from the left and right of 3 embryos) for each of the anterior, middle and posterior domains of the developing cup. The origin of the tracks has been arbitrarily set to the intersection of the X/Y axis and the early (coloured) and late (Grey) phases of migration have been highlighted.

(**B**) Mean tracks for anterior EON of control (Black), *cxcr4b*^*t26035*^ (Blue) and *cxcl12a*^*t30516*^ (Green) embryos.

**Figure S6.** Morphogenetic movements of skin cells in *cxcr4b*^*t26035*^ and *cxcl12a*^*t30516*^ embryos.

Tracks showing migration of skin cells of control (Black), *cxcr4b*^*t26035*^ (Blue) and *cxcl12a*^*t30516*^ (Green) embryos. 12 tracks are represented for cells overlying the anterior, middle and posterior domains of the developing cup. The origin of the tracks has been arbitrarily set to the intersection of the X/Y axis and the early (coloured) and late (Grey) phases of migration have been highlighted. Morphogenetic movements of individual skin cells showed no obvious differences in control versus *cxcr4b*^*t26035*^ or *cxcl12a*^*t30516*^ embryos.

**Figure S7.** Crispr-Cas9 deletion of the Neurog1-regulated CRM upstream of the *cxcr4b* locus.

(**A**) Schematic representation of the *cxcr4b* locus indicating the position of exons of the *cxcr4b* gene (orange) and E-box clusters, which are color-coded depending on the nature of the E-box sequences. The position of the genomic sequences found in the *TgFOS(cxcr4b:eGFP)* transgene and a schematic representation of the PCR fragment used to genotype potential deletions with the sequence of the E-box cluster are also shown.

(**B**) Ethidium bromide stained agarose gel showing the results of PCR genotyping for the induction of deletions of the CAGATG E-box cluster. The magenta arrowhead shows the 500bp control band. A 200bp band appears (black arrowhead) when the sgRNA pair is injected with Cas9 but not when the sgRNA pair or Cas9 is injected alone (bp, base pair).

## Supplemental Movies

**Movie S1.** Time-laps imaging reveals a convergence migration of EON towards the centre of the future olfactory cup.

Movie showing an acquisition series of *Tg(−8.4neurog1:gfp)* labelled EON in a control embryo. The movie shows the entire acquisition from 12 hpf to 27 hpf with frames every 7 min.

**Movie S2.** Olfactory cups in *neurog1*^*hi1059*^ mutants exhibit delays in convergence.

Movie showing an acquisition series of *Tg(−8.4neurog1:gfp)* labelled EON in a *neurog1*^*hi1059*^ mutant embryo. The movie shows the whole acquisition from 12 hpf to 27 hpf with frames every 7 min.

**Movie S3.** Tracking EON behaviour at single-cell resolution.

Behaviour of EON located in the anterior, middle and posterior thirds of the neurog1:GFP+ population was established by manually tracking H2B-RFP positive nuclei. The movie is divided into 6 parts representing: the acquisition showing neurog1:GFP+ cells (Green) and their nuclei (Magenta), tracking of anterior, middle and posterior EON cells, all tracking combined and a still image showing the tracks from anterior, middle and posterior cells.

**Movie S4.** Tracking EON behaviour at single-cell resolution in *neurog1*^*hi1059*^ mutant embryo.

The movie is divided into 6 parts representing: the acquisition showing neurog1:GFP cells, tracking of anterior, middle and posterior EON cells, all tracking combined and a still image showing the tracks from anterior, middle and posterior cells; the early phase of migration that is affected in the mutant is highlighted (Magenta).

**Movie S5.** Tracking EON behaviour at single-cell resolution in *cxcr4b*^*t26035*^ mutant embryo.

The movie is divided into 3 parts representing: the acquisition showing neurog1:GFP cells, tracking of a pair of anterior EON cells and a still showing the anterior tracks; the early phase of migration that is affected in the mutant is highlighted (Blue).

**Movie S6.** Tracking EON behaviour at single-cell resolution in *cxcl12a*^*t30516*^ mutant embryo.

The movie is divided into 3 parts representing: the acquisition showing neurog1:GFP cells, tracking of a pair of anterior EON cells and a still showing the anterior tracks; the early phase of migration that is affected in the mutant is highlighted (Green).

**Movie S7.** Re-expressing Cxcr4b in *neurog1*^*hi1059*^ mutant embryo rescues anterior EON behaviour.

The movie is divided into 3 parts representing: the acquisition showing neurog1:GFP cells, tracking of an anterior EON cell and a still showing the anterior track; the early phase of migration is highlighted (Cyan).

**Movie S8.** Rescue transgene expression does not affect anterior EON behaviour in a control context.

The movie is divided into 3 parts representing: the acquisition showing neurog1:GFP cells, tracking of an anterior EON cell and a still showing the anterior track; the early phase of migration is highlighted (Light Blue).

