## Supplementary figures and images for "Morphogenesis is transcriptionally coupled to neurogenesis during peripheral olfactory organ development"

### Supplementary Figure S1

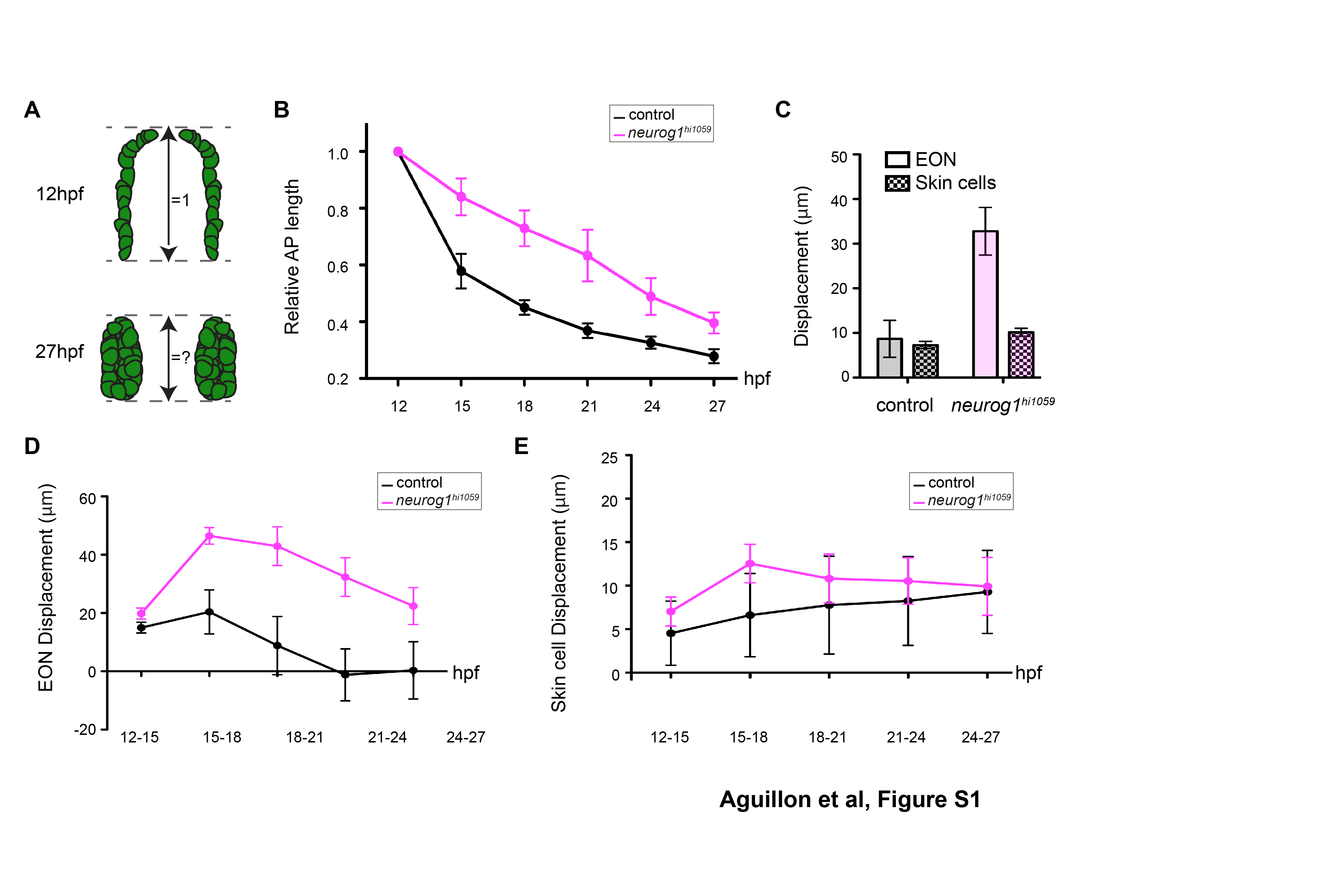

### Supplementary Figure S2

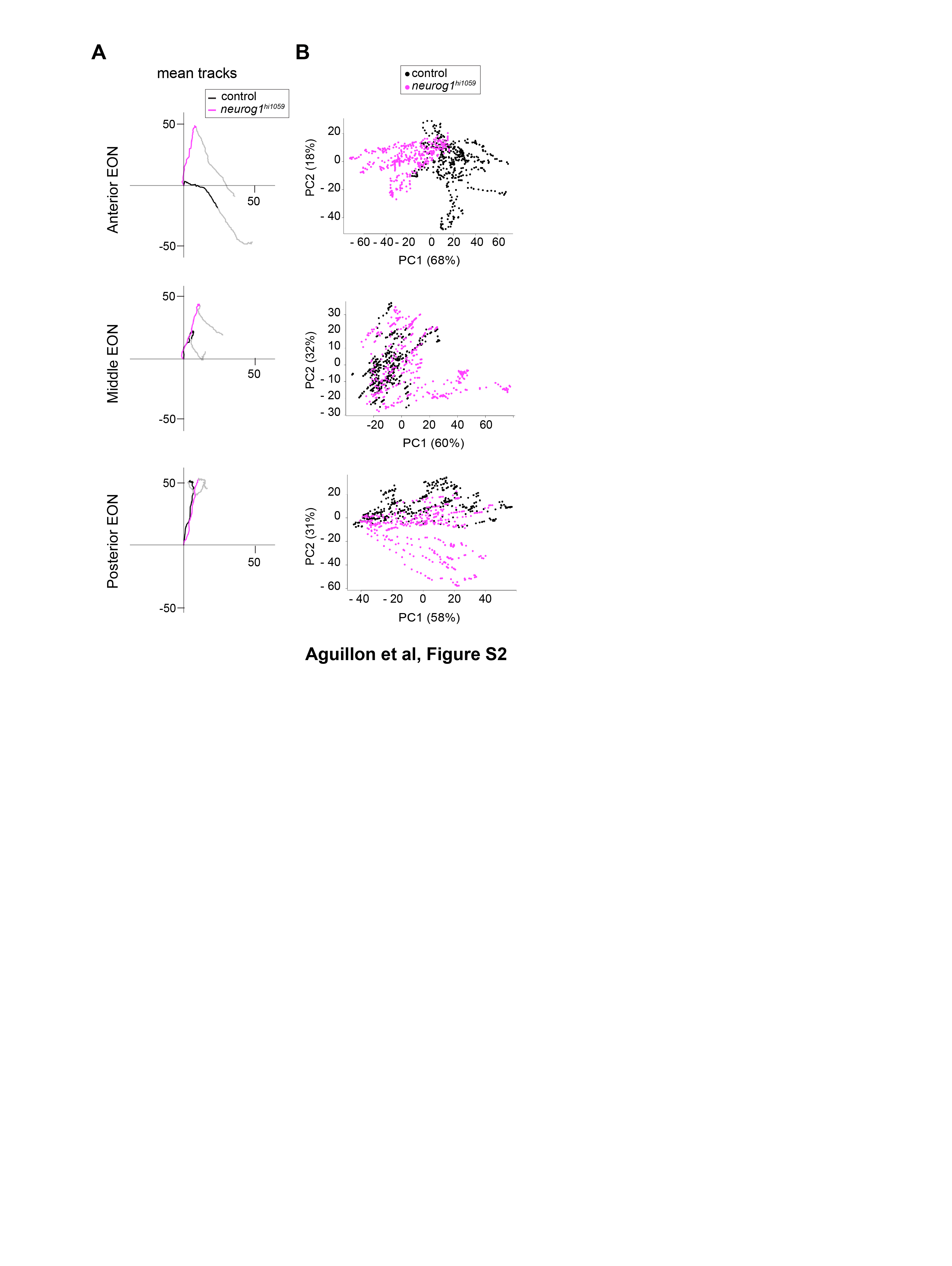

### Supplementary Figure S3

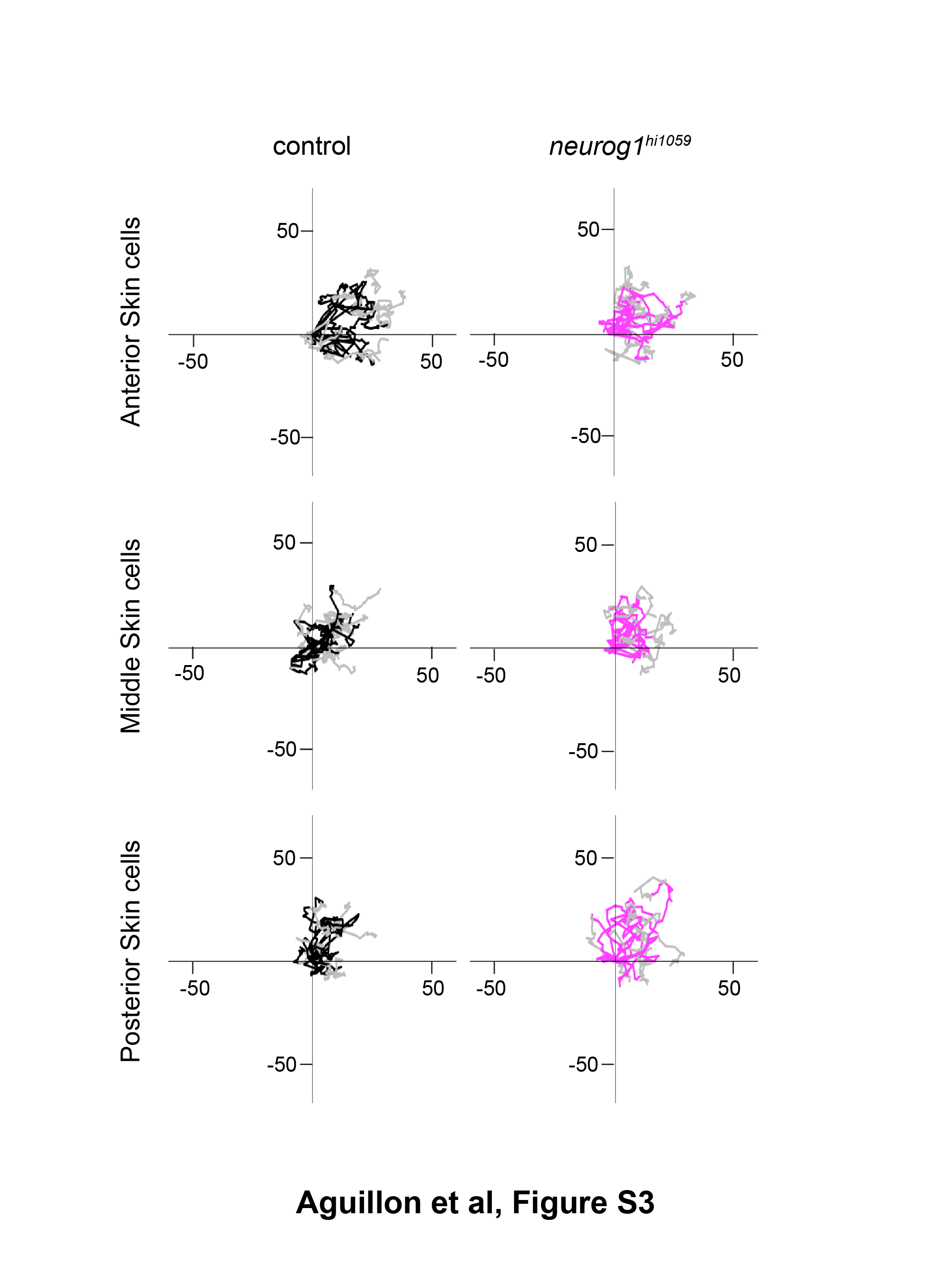

### Supplementary Figure S4

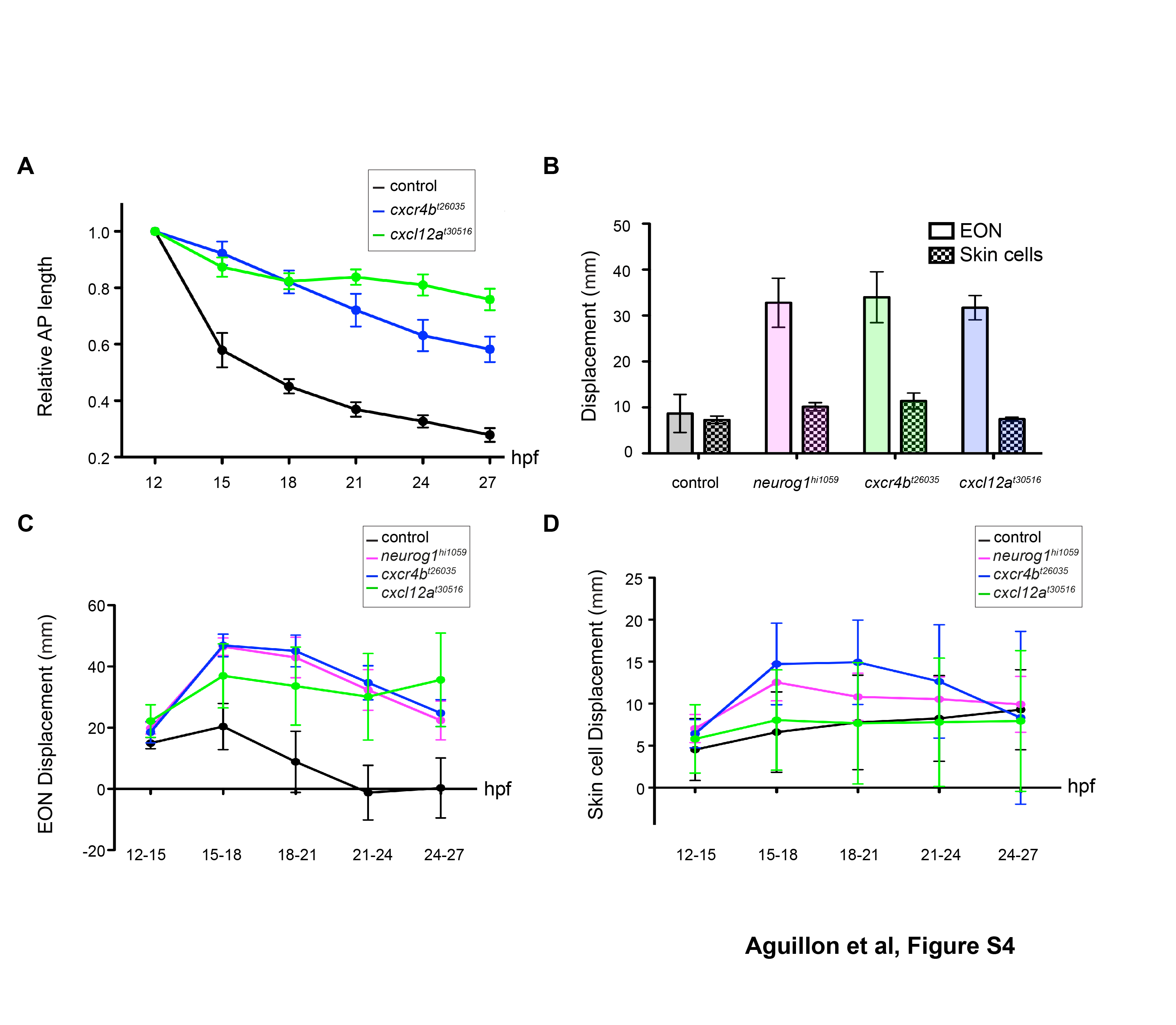

### Supplementary Figure S5

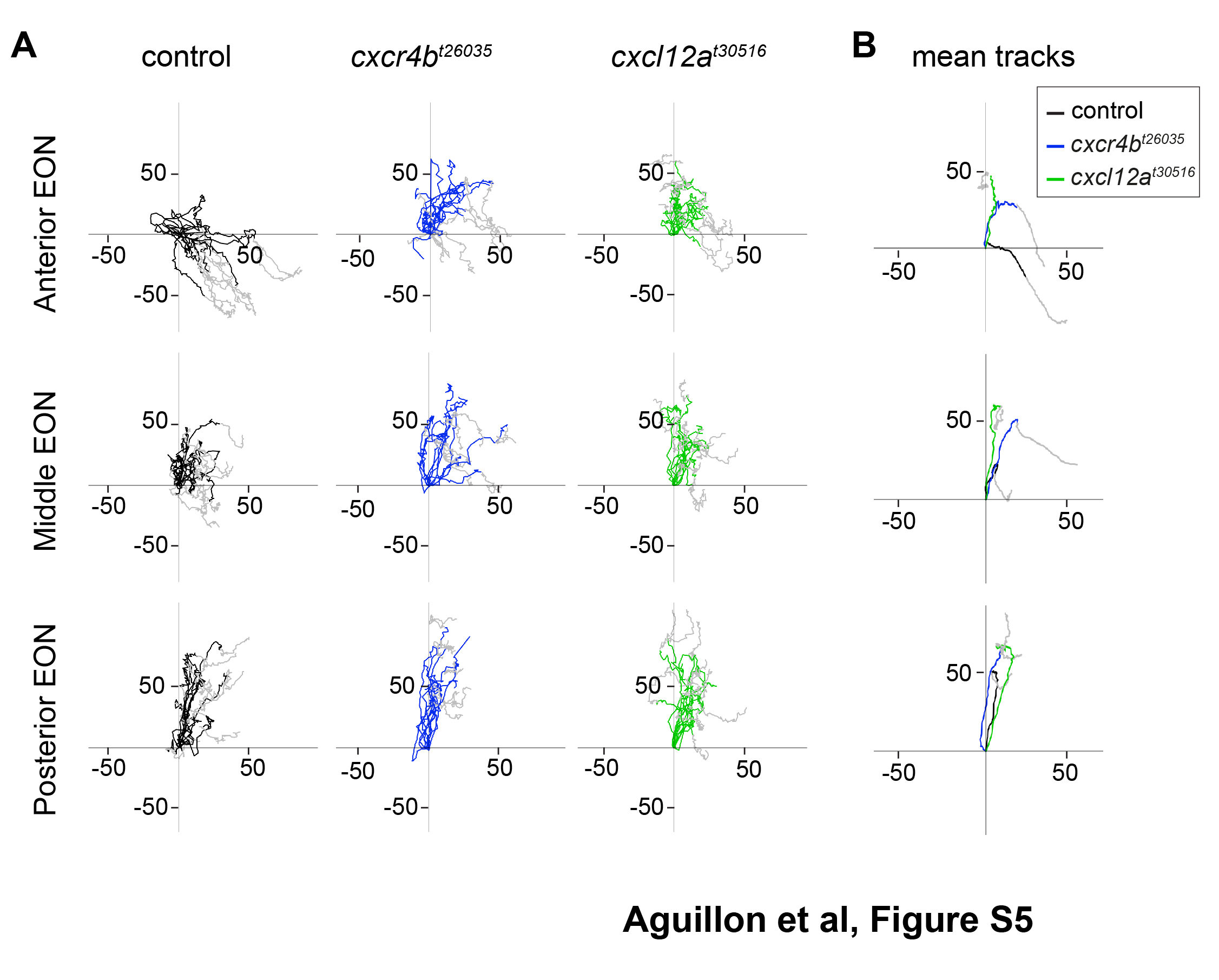

### Supplementary Figure S6

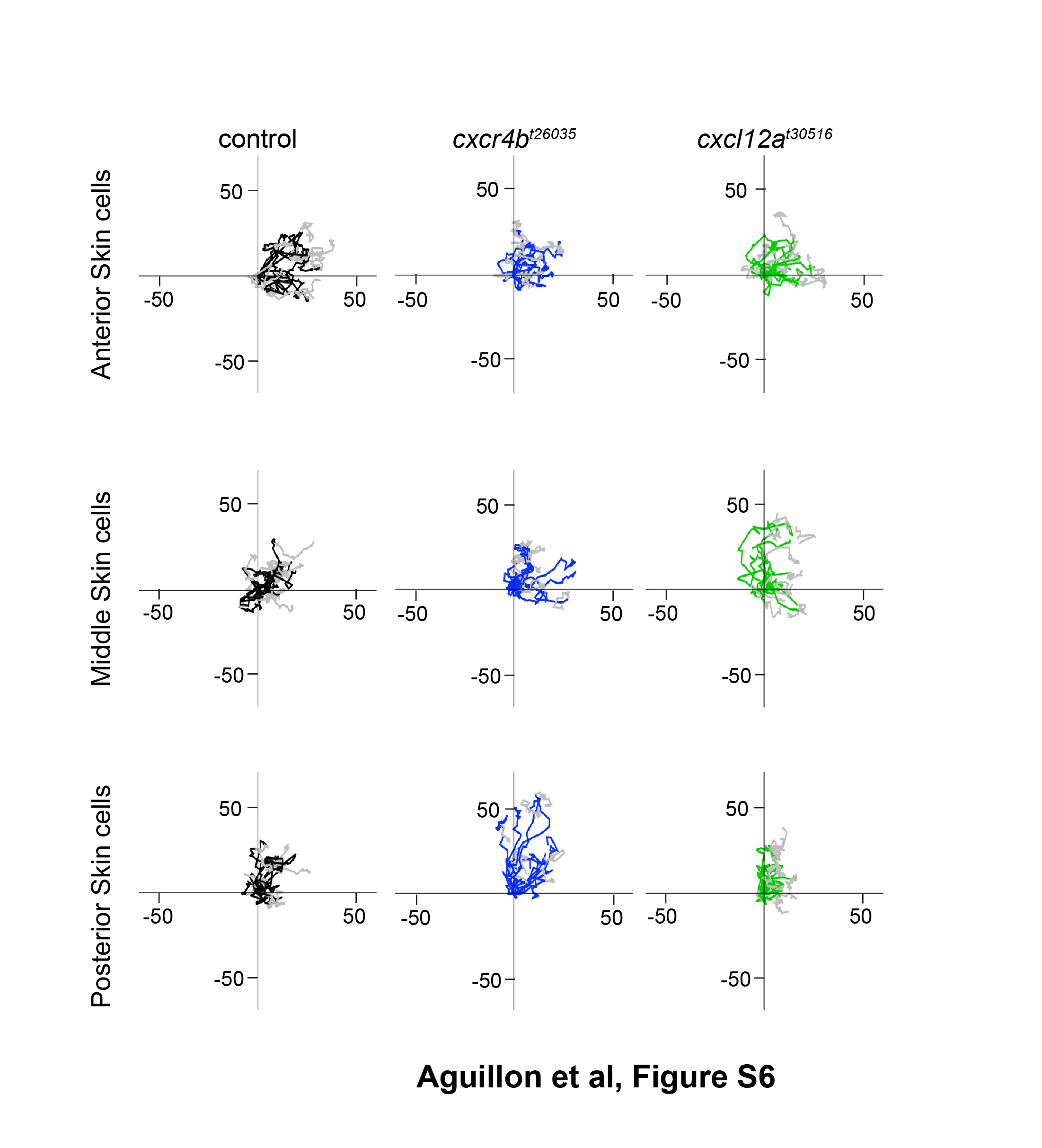

### Supplementary Figure S7

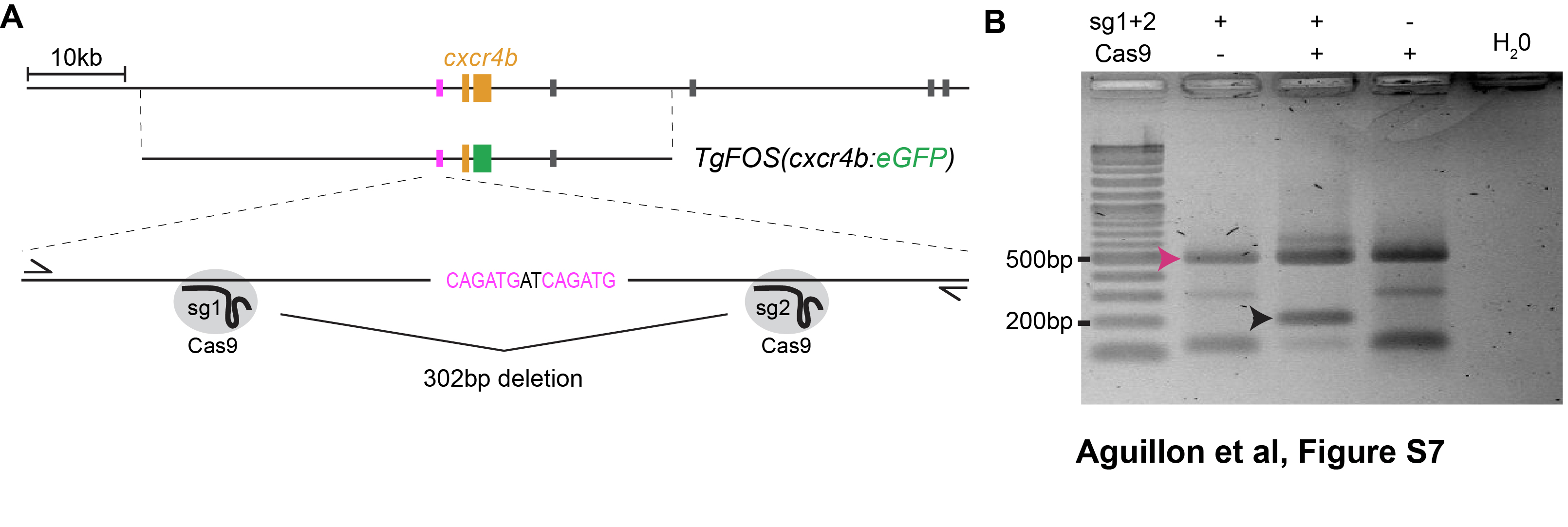
